# Long non-coding RNAs bind to proteins relevant to the ethanol tolerance in yeast: a systems biology view

**DOI:** 10.1101/2021.02.07.430053

**Authors:** Lucas Farinazzo Marques, Ivan Rodrigo Wolf, Lucas Cardoso Lazari, Lauana Fogaça de Almeida, Amanda Piveta Schnepper, Luiz Henrique Cardoso, Leonardo Nazário de Moraes, Rejane Maria Tommasini Grotto, Rafael Plana Simões, Érica Ramos, Guilherme Targino Valente

## Abstract

The ethanol disturbs the cell cycle, transcription, translation, protein folding, cell wall, membranes, and many *Saccharomyces cerevisiae* metabolic processes. Long non-coding RNAs (lncRNAs) are regulatory molecules binding onto the genome or proteins. The number of lncRNAs described for yeast is still scarce, and little is known concerning their roles in the system. There is a lack of knowledge concerning how lncRNAs are responsive to the ethanol tolerance in yeast and whether they act in this tolerance. Hence, by using RNA-Seq data from *S. cerevisiae* strains with different ethanol tolerance phenotypes, we found the severe ethanol responsive lncRNAs. We modeled how they participate in the ethanol tolerance by analyzing lncRNA-protein interactions. The results showed that the EtOH tolerance responsive lncRNAs, in both higher tolerant and lower tolerant phenotypes, work on different pathways: cell wall, cell cycle, growth, longevity, cell surveillance, ribosome biogenesis, intracellular transport, trehalose metabolism, transcription, and nutrient shifts. In summary, lncRNAs seems to interconnect essential systems’ modules to overcome the ethanol stress. Finally, here we also found the most extensive catalog of lncRNAs in yeast.

## Introduction

Biofuels can be produced from different sources (Demirbas 2017). Bioethanol is the most important biofuel as the most promising gasoline substitute (Chakraborty *et al*. 2012; Gupta and Verma 2015). The yeast *Saccharomyces cerevisiae* is the most used organism for bioethanol production (Mussatto *et al*. 2010; Demeke *et al*. 2013). Thus, understanding relevant factors to improve ethanol yield is essential for an evergrowing environment that increasingly demands more fuel.

However, increasing the ethanol (EtOH) concentration compromises the yeast survival affecting its growth rate and production (Stanley *et al*. 2010; Auesukaree 2017). For instance, EtOH rapidly affects the yeast’s plasma membrane integrity (Ding *et al*. 2009; Ma and Liu 2010; Navarro-Tapia *et al*. 2016), leading to protein dysfunction and denaturation, affecting the molecules intake (e.g., glucose and amino acids), and causing an efflux of nucleotides and potassium (Ding *et al*. 2009; Ma and Liu 2010).

Experiments overexpressing or repressing candidate genes change the EtOH tolerance in yeast (Alper *et al*. 2006; Teixeira *et al*. 2009; Mussatto *et al*. 2010; Lewis *et al*. 2010; Swinnen *et al*. 2012). However, the complex molecular mechanisms concerning this pathway are still poorly understood.

The long non-coding RNAs (lncRNAs) promptly respond to external stimuli (Yamashita *et al*. 2016), regulating the gene expression and epigenetic modifications (Anderson *et al*. 2015). Furthermore, the few known cases concerning lncRNA-protein interactions suggest that lncRNAs work as a framework for macromolecular complexes assembly, or bait transcription factors dampening the association of these proteins to the DNA, or guide the chromatin modifiers (Tripathi *et al*. 2010; Geisler and Coller 2013; Ferrè *et al*. 2016; Li *et al*. 2019).

The lncRNA description for *S. cerevisiae* is scarce. Only 18 lncRNAs are appropriately described and annotated for this species (Till *et al*. 2018). These lncRNAs are directly involved in metabolic changes, sexual differentiation initiation, and other unknown processes (Yamashita *et al*. 2016). Recently, experiments showed that four ncRNAs affect the transcriptional systems as a whole mediated by a *trans* effect on transcription factors. These ncRNAs may be associated with the EtOH tolerance and stress response (Balarezo-Cisneros *et al*. 2020). However, little is known concerning the roles of lncRNA-protein interactions for all species.

Here we aimed to raise hypotheses concerning the relationship between lncRNAs and the EtOH tolerance. For this purpose, we focused on analyzing the interactions between the EtOH tolerance responsive lncRNAs and proteins. RNA-Seq data from *S. cerevisiae* strains with different EtOH tolerance phenotypes (higher and lower EtOH tolerant ones) were used to seek the lncRNAs expressed during the severe EtOH stress; in this case, we developed a pipeline to assemble these lncRNAs. Then, we analyzed how the lncRNAs impact the EtOH tolerance based on the prediction of lncRNA-protein interactions, guilt-by-association, and information flow throughout network approaches. The main pathways that EtOH stress-responsive LncRNAs work on are the cell wall, cell cycle and growth, cell longevity, cell surveillance, ribosome biogenesis, intracellular transport, trehalose metabolism, transcription, and nutrients shifts. Our findings indicate that EtOH stress-responsive lncRNAs interconnect essential systems’ modules in a sort of strain-specific way to surpass a stressed environment’s challenges. Altogether, here we provide insights and hypotheses on how lncRNAs may be working to the EtOH tolerance.

## Material and Methods

### Strains selection, ethanol tolerance definitions, and sequencing

The haploid strains BMA64-1A (Euroscarf/20000A), BY4742 (SGD/BY4742), X2180-1A (SGD/X2180-1A), BY4741 (SGD/BY4741), SEY6210 (SGD/SEY6210), and S288C (SGD/S288c) were obtained from Euroscarf (European *Saccharomyces cerevisiae* Archive for Functional Analysis) or NRBP (National Bioresources Project).

For the EtOH tolerance experiments, strains were grown overnight in YPD (2% of peptone, 1% of yeast extract, and 2% of glucose) and further diluted to an OD_600_ of 0.2. Then, 100 μL of cells were harvested by centrifugation at 2,000 RPM at 4°C for 5 min. Pellets were resuspended using YPD with different EtOH or physiological solution concentrations (the treatment and control condition, respectively) in plate-wells. Plates were incubated at 30°C for 1h and shaken at 120 RPM; the EtOH or physiological solution ranged from 2% to 32% (v/v). The content of each plate-well was plated on YPD and incubated at 30°C. Visual inspection allowed to set up the highest EtOH tolerance level supported for each strain.

Total RNA of cells under control and treatment conditions (the highest EtOH level supported per strain) was extracted from 1.5 mL of cells using Trizol after a cell wall digestion using Lyticase. The RNA quality was checked by electrophoresis in agarose gel and Bioanalyzer, and Nanodrop and Qubit estimated the RNA quantification. The samples were treated with TURBO DNase (ThermoFisher), and rRNA depletion was performed. The transcriptome was obtained for 36 samples (6 strains x 2 (treatment and control) x 3 replicates). The genomic DNA of BMA64-1A was extracted using phenol-chloroform, and the quality and quantity were estimated using the equipment mentioned.

The RNA-Seq was performed by the LcScience (Texas, USA) Company using the Illumina HiSeq 4000 (100 nt, paired-end reads, insert size of 24-324 bp, and at least 40 million reads per sample); the company ensured the absence of small RNAs (<200 nts). The GenOne (Rio de Janeiro, Brazil) sequenced the genome using the Illumina HiSeq 2500 (250 nt, paired-end reads, 1 Gb of throughput, and insert size ~500 bp).

### Genome assembling, lncRNA identification, and differential expression

The paired-end reads of the BMA64-1A genome were cleaned with Trimmomatic v.0.36 and independently assembled using AbySS v.2.0.2 (Jackman *et al*. 2017), IDBA v.1 (Peng *et al*. 2010), MIRA v.4.0.2, SPAdes v.3.10.1 (Bankevich *et al*. 2012) and Velvet v.1.2.10 (Zerbino and Birney 2008), varying parameters and assembling strategies. For an individual chromosome assembling, the reads were mapped against the reference genome (S288C version R64-2-1) using the HISAT2 v.2.1.0 (Kim *et al*. 2015), and the mapped reads were assembled using IDBA v.1 (Peng *et al*. 2010).

The genomic assembling metrics of each assembly were obtained with QUAST v. 4.5 (Gurevich *et al*. 2013). The QUAST values were normalized, generating a score from 0 to 1. Since QUAST generate many metrics for the assembling and reference, the score mentioned was calculated for each assembling considering all QUAST-normalized metrics (equation 1); the assembling with score value ≈0 was considered the most similar to the reference genome, and we assumed as the final data:

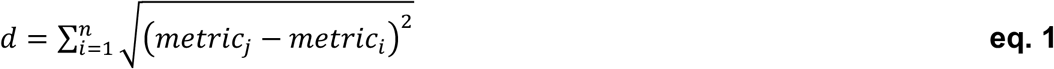

where *metric_j_* and *metric_i_* are the metrics of the reference genome and a given assembling, respectively. To obtain *d*, all *n* metrics were considered.

For the BMA64-1A genome annotation, the transcripts of S288C from SGD were mapped over the best assembling using GMAP (Wu and Watanabe 2005), and a *de novo* annotation was performed using MAKER v.2.32 (Cantarel *et al*. 2008). A manual curation comparing the two annotations was performed: 1-annotations present in both strategies were considered correct; 2-by visual inspection of the reference genome, the regions with disagreements passed by manual adjustments. Finally, we adjusted the annotations using the results from AGAPE (Song *et al*. 2015). The protein sequences were translated from annotated regions for further analysis.

The paired-end reads for all RNA-Seq libraries were trimmed using Trimmomatic v.0.36 (Bolger *et al*. 2014) and used to identify the lncRNAs.

We developed a pipeline to assembly the lncRNAs. Overall, the filtered reads were mapped over coding sequences of many different species. After, the non-mapped paired-reads were assembled using different algorithms, and a score was calculated to rank the best assembling. The transcript redundancies within strains were excluded, and the remaining transcripts were mapped over the genome of each strain to exclude spurious assembling. The mapped transcripts that may be coding sequences or mobile elements were excluded, and the remaining sequences were rechecked concerning their potential coding (Figure 1). Details of this pipeline are below described.

**Figure 1:**
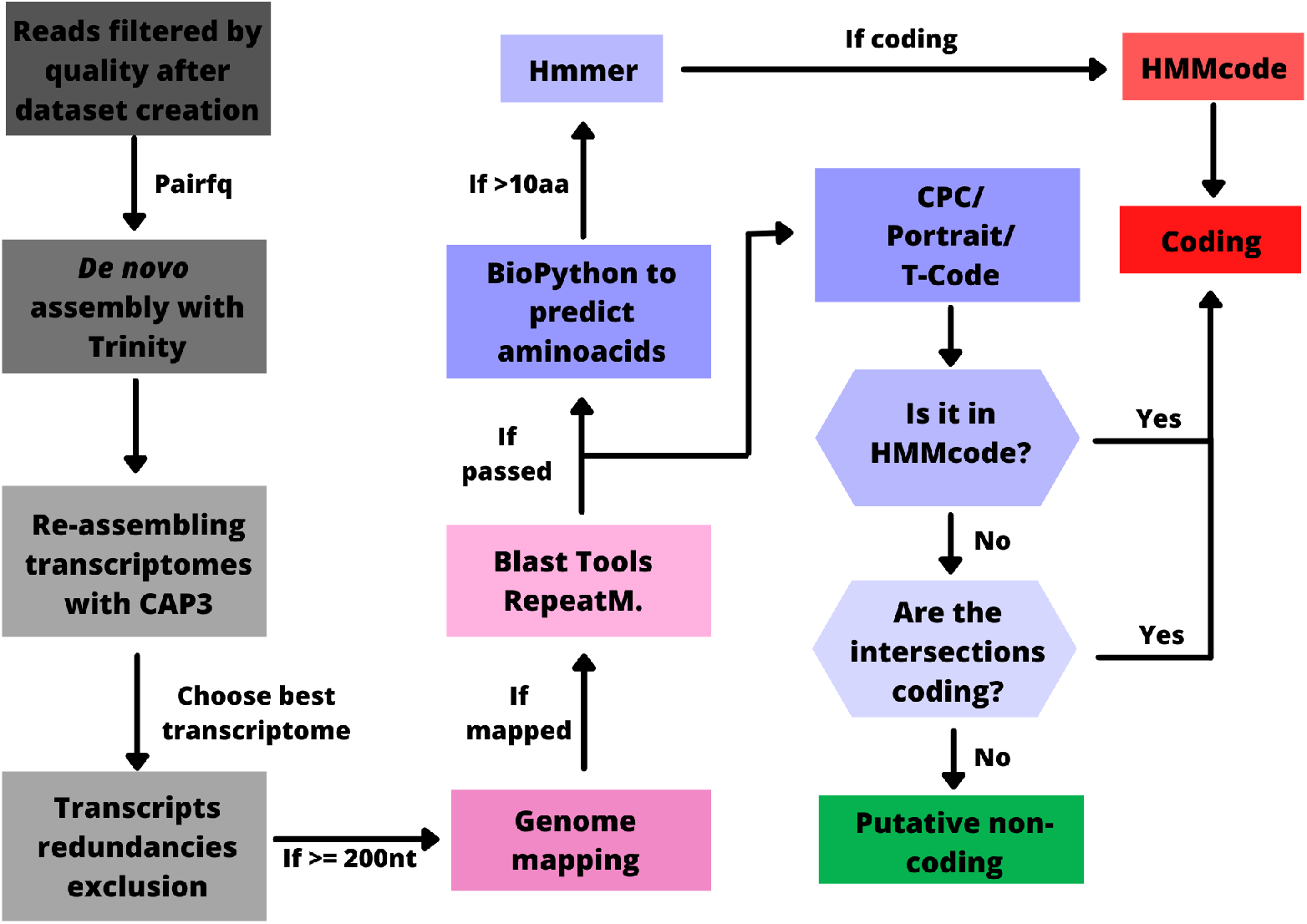
Pipeline to assembly the lncRNAs.

It was created a dataset with millions of sequences including: 1-coding sequences (CDS), and proteomes of eukaryotes and bacteria; 2-genomes; 3-microRNAs precursors; 4-ncRNA families present in the Rfam database (Burge *et al*. 2013), being the lncRNAs (Rfam accession number 01884) the only exception; 5-mobile elements. The RNA-Seq filtered reads of each strain were independently aligned on nucleotide sequences of the database mentioned using HISAT2 v.2.1.0 (Kim *et al*. 2015). Then, we assembled the non-aligned read-pairs selected by Pairfq script (The MIT License) since the ones are reads without similarity with coding, ncRNAs (excepting lncRNAs), mobile elements, and mitochondrial and contaminants genomes; reads that lost a member of the pair were excluded of this assembling (Figure 1).

The “Single Assembler Multiple Parameters” strategy (He *et al*. 2015) was used for the *de novo* assembling mentioned; it was performed parameter adjustments for each step using the S288C reads, and then, we applied the best parameter set to independently assembly the reads of all other strains. First, the Velvet/Oases (Zerbino and Birney 2008; Schulz *et al*. 2012), Trinity (Haas *et al*. 2013), IDBA-tran (Peng *et al*. 2012), and rnaSPAdes (Bankevich *et al*. 2012) were independently tested to determine which one was the best assembler for our dataset. For Velvet/Oases, rnaSPAdes, and IDBA-tran parameters, the *kmers* ranged from 19 to 81, and for Velvet/Oases and rnaSPAdes, we had set-up an automatic coverage cutoff and no scaffolding assembling. For Trinity assembler, we set-up the *kmers* ranging from 19 to 31. These programs were set-up to assemble only transcripts ≥200 nts when this option was available (Figure 1).

All assembled transcriptomes within each assembler were merged in a single file, sequences with ≥10% of “Ns” (non-identified nucleotides) were removed, followed by a generation of 1 transcriptome per assembler; this re-assembling was performed using CAP3 (Huang 1999). The singlets and contigs from CAP3 were merged, and the filtered reads (the ones used in the first assembling) were mapped over this file using Bowtie2 (Langmead and Salzberg 2012). A score was used (equation 2) to evaluate the final assembling, and the highest value indicates the best transcriptome. As mentioned, after these adjustments of the *de novo* assembling pipeline using the reads of S288C, the same settings were applied to independently assemble the lncRNAs of other strains (Figure 1).

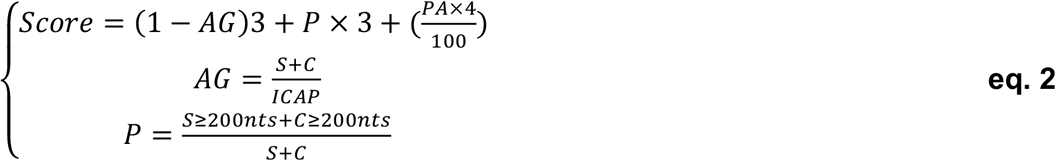

where *AG* (the assembling gain) is a rate of how many sequences the CAP3 used to assemble. The *P* is the percentage of CAP3 assembled sequences ≥200 nts, and the *PA* is the percentage of reads aligned using Bowtie2 over CAP3 assembling. The *S* and *C* are the numbers of sequences in the “singlets” and “contigs” CAP3’s output, respectively. The *ICAP* is the number of sequences with ≤10% of “Ns” (assembled by IDBA, rnaSPAdes, Trinity, or Velvet/Oases) used as CAP3 input. The *PA* and *P* range from 0 to 100, while *AG* ranges from 0 to 1. The *AG* rates ≈0 are considered better values since it expresses that a higher number of input sequences was re-assembled by CAP3. The *P* and *PA* ≈1 are better, expressing that CAP3 assembled a higher number of large transcripts, and a higher number of reads was used by the first assemblers (the ones before CAP3), respectively.

The set of non-redundant transcripts found using CD-HIT (Li and Godzik 2006) were merged. The non-redundant sequences of each strain were mapped over its own genomes using GMAP (Wu and Watanabe 2005). The mapped transcripts were then rechecked concerning their potential to be coding, ncRNAs (except lncRNAs), repetitive elements, or contaminant genomes (viruses, bacteria, and mitochondria). For this purpose, the mapped transcripts were: 1-aligned against the proteomes using Blastx; 4-aligned against bacteria, viruses, and mitochondrial genomes using GMAP (Wu and Watanabe 2005). The data distribution of all results was analyzed to establish cutoffs to filter out undesirable sequences (Figure 1).

To verify the lack of coding regions on filtered sequence transcripts were protein translated (≥10 aa) using the Getorf (Rice *et al*. 2000), and transcripts/proteins were evaluated using Hmmer (Mistry *et al*. 2013) with Pfam database v.31.0 (Finn *et al*. 2014), Tcode (Rice *et al*. 2000), Portrait (Arrial *et al*. 2009) and CPC (Kong *et al*. 2007). The transcripts with motifs fitting Hmm models using Hmmer were assumed as potentially coding, whereas the other sequences that fit coding sequences using at least two out of the other three programs (Tcode, Portrait, or CPC) were defined as coding sequences. Hence, sequences not found neither by Hmmer nor by two other programs were assumed as putative lncRNAs (Figure 1).

The RNA-Seq filtered reads of each strain were mapped over its genomes using HISAT2 v.2.10 (Kim *et al*. 2015) (“--no-softclip”), the counting per transcript was obtained using the Bedtools multiBamCov v.2.26.0 (Quinlan and Hall 2010), and the lncRNAs differentially expressed comparing treatment *vs*. control was provided by DESeq2 (Love *et al*. 2014) considering a false discovery rate <0.01.

### LncRNA-protein interaction prediction and analysis

It was predicted the lncRNAs-protein interactions (LNCPI), considering proteins ≥32 amino acids of each strain (downloaded from SGD and translated from the BMA64-1A genome assembling), using the lncPRO tool (Lu *et al*. 2013). The probability distributions of interactions follow a normal distribution, then the ones ≥0.95 of probability were selected for further analysis.

We downloaded yeast’s protein-protein interactions (PPI) from Biogrid (Chatr-Aryamontri *et al*. 2013) and MINT (Licata *et al*. 2012), gene regulatory networks from the YTRP database (Yang *et al*. 2014), and metabolic networks from REACTOME (Fabregat *et al*. 2014). Only physical interactions among proteins were selected from Biogrid, and redundancies with MINT were excluded; we excluded the interactions present in the NEGATOME (Blohm *et al*. 2014). The networks were combined to create a single integrated network (Uninet). The Uninet was unified with the LNCPI of each strain generating 6 networks (one per strain and each of them including its LNCPI strain-specific) (equation 3). These networks were considered undirected graphs.

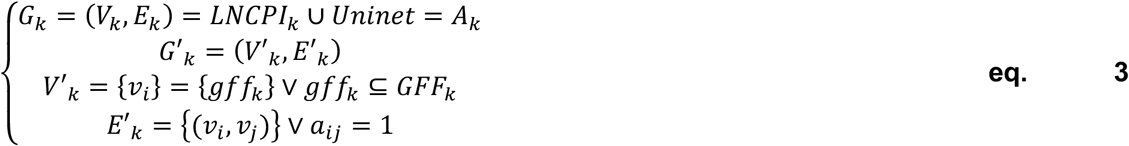

which *G_k_* is an adjacency matrix *A_k_* (where *a_ij_* is an element of *A_k_*) of a given strain (*k*). The *G’_k_* is a subgraph from *G_k_*. *V’_k_* is a subset of nodes *v_i_* from *G’_k_* generated based on GFF annotation (*gff_k_*). *E’_k_* is a subset of edges of *V’_k_* from *G’_k_*. Hence, *G’_k_* are strain-specific networks harboring their respective LNCPI as well.

To assess the influence of EtOH on the lncRNA target-protein and how the lncRNAs could help to the EtOH tolerance, we analyzed the information flow throughout LNCPI networks from each differentially expressed lncRNA (hereafter referred to as lncRNA-propagation analysis). For this analysis, we selected the up-regulated edges (log2 fold-change ≥ 1E-6) of each strain (the network is the *G’_k_* depicted in equation 3). Unconnected nodes or nodes with self-loops were excluded. The network propagation using the diffusion algorithm (Cowen *et al*. 2017) was assessed starting from all differential expressed lncRNAs. After, the lncRNAs and their first neighbors were selected, and only the top 20 ranked nodes from diffusion information were retrieved to compose new subgraphs (the nodes with self-loops or unconnected nodes were excluded again). Altogether, we could assess the participation of lncRNAs in the EtOH tolerance and whether this compound triggers similar effects on systems harboring different lncRNAs and interactions. The same processes were performed for lncRNAs and genes down-regulated (log2 fold-change ≤-1E-6).

The gene ontology of lncRNA target-proteins was analyzed using g:Profiler (Reimand *et al*. 2016). The enriched terms were summarized by REVIGO (Supek *et al*. 2011). For this purpose, we considered all target-proteins indiscriminate neither by differential expression nor by strain.

## Results

The highest EtOH tolerance level observed for each strain allowed classifying the ones as higher tolerant (HT) or lower tolerant (LT) phenotypes. The HT strains are BMA64-1A (tolerates 30% of EtOH), BY4742 (tolerates 26% of EtOH), and X2180-1A (tolerates 24% of EtOH). The strains BY4741 (tolerates 22% of EtOH), SEY6210 (tolerates 20% of EtOH), and S288C (tolerates 20% of EtOH) were considered LTs.

A total of 6,410,152 filtered reads with ~250 nts length allowed to achieve 196 assemblings of the BMA64-1A genome (Table 1). The best assembling has 11,880,801 bp, with 683 scaffolds, and with 134.88X of coverage. Only 36 out of 6,551 annotated coding genes did not present any similarity with the S288C genome (the reference genome).

**Table 1:** Metrics of BMA64-1A genome considering all assembling. *: metrics of selected assembling; **AS:** assembling score.

| Value | N50 | Contigs/scaffolds | GC% | Genome size | Genes | AS |
| --- | --- | --- | --- | --- | --- | --- |
| Minimum | 1,437 | 17 | 37.93 | 126,580 | 1,744 | 0.054 |
| Maximum | 924,585 | 27,639 | 38.34 | 12,162,499 | 5,872 | 0.63 |
| *Selected | 202,259 | 683 | 38.06 | 11,880,801 | 5,668 | 0.29 |

The best lncRNA assemblings pipeline uses the Trinity, followed by a reassembling using CAP3. Although Velvet/Oases outperforms the Trinity (see the score in Table 2), the latter presented the highest computational performance. Hence, all lncRNA assembling used Trinity giving from 87 to 259 lncRNAs identified after the filtering steps (Table 3).

**Table 2:**
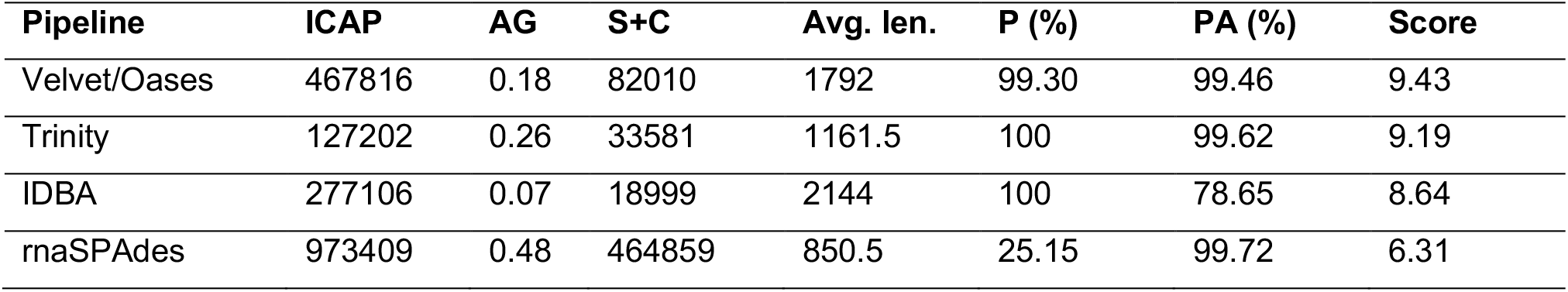
Overview of *de novo* final assemblings of S288C’s lncRNAs. The ICAP, AG, S+C, P, PA, and Score are described in equation 2. **Avg. len.:** the average sequence length of CAP3 re-assembling.

**Table 3:**
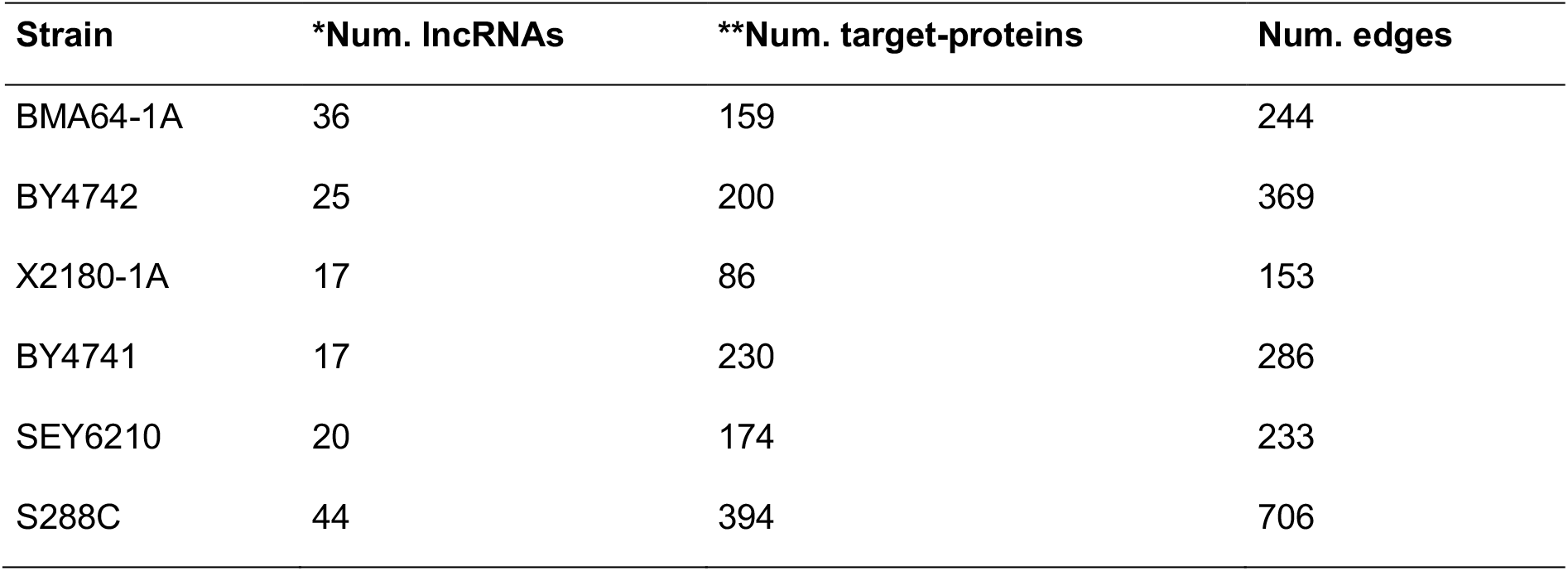
Summary of LNCPI. *: the number of lncRNAs with at least one interaction; **: the number of proteins potentially binding to lncRNAs.

| Strain | *Num. lncRNAs | **Num. target-proteins | Num. edges |
| --- | --- | --- | --- |
| BMA64-1A | 36 | 159 | 244 |
| BY4742 | 25 | 200 | 369 |
| X2180-1A | 17 | 86 | 153 |
| BY4741 | 17 | 230 | 286 |
| SEY6210 | 20 | 174 | 233 |
| S288C | 44 | 394 | 706 |

Most lncRNAs range from ~200-400 nts (Figure 2), being the lncRNA transcr_28768 of S288c the largest one (2,739 nts). Interestingly, all lncRNAs do not present similarities to those previously published (Parker *et al*. 2017, 2018).

**Figure 2:**
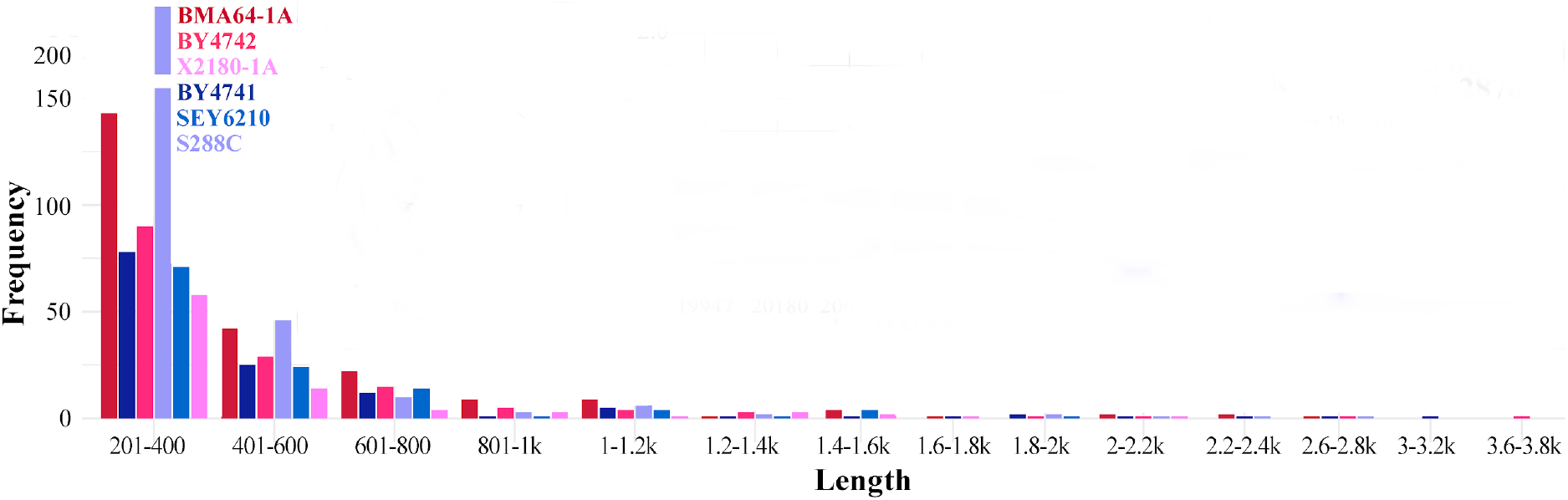
The distribution of lncRNA lengths.

The filtered LNCPI networks (probability ≥95%) harbor most of the identified lncRNAs. The filtered networks presented from 17 to 44 lncRNAs and from 86 to 394 target-proteins. The number of edges found is higher than the number of target-proteins, indicating that some lncRNAs interact with multiple proteins (Table 3). Indeed, the average of target proteins divided by the number of lncRNAs is ~8.1.

The EtOH stress-responsive lncRNAs lie in four functional categories. These categories were identified based on the function of target-proteins selected by lncRNA-propagation analysis from each differentially expressed lncRNAs. The 1^st^ category (“essential process”) includes genes related to transcription, replication, and ribosome biogenesis. The 2^nd^ (“membrane dependent process”) includes genes related to signaling and cell division, membranes and cell wall, and intracellular transport. The 3^rd^ (“metabolic process”) includes genes related to oxidative stress response, diauxic shift, fermentation, and other metabolisms. The 4^th^ includes genes of the “degradation process” (Figure 3A). Moreover, the whole set of target-proteins (without considering differential expression and strain-specificities) are mainly related to “RNA polymerase assembling” and “negative regulation of protein kinase activity by protein phosphorylation” (Figure 3B).

**Figure 3:**
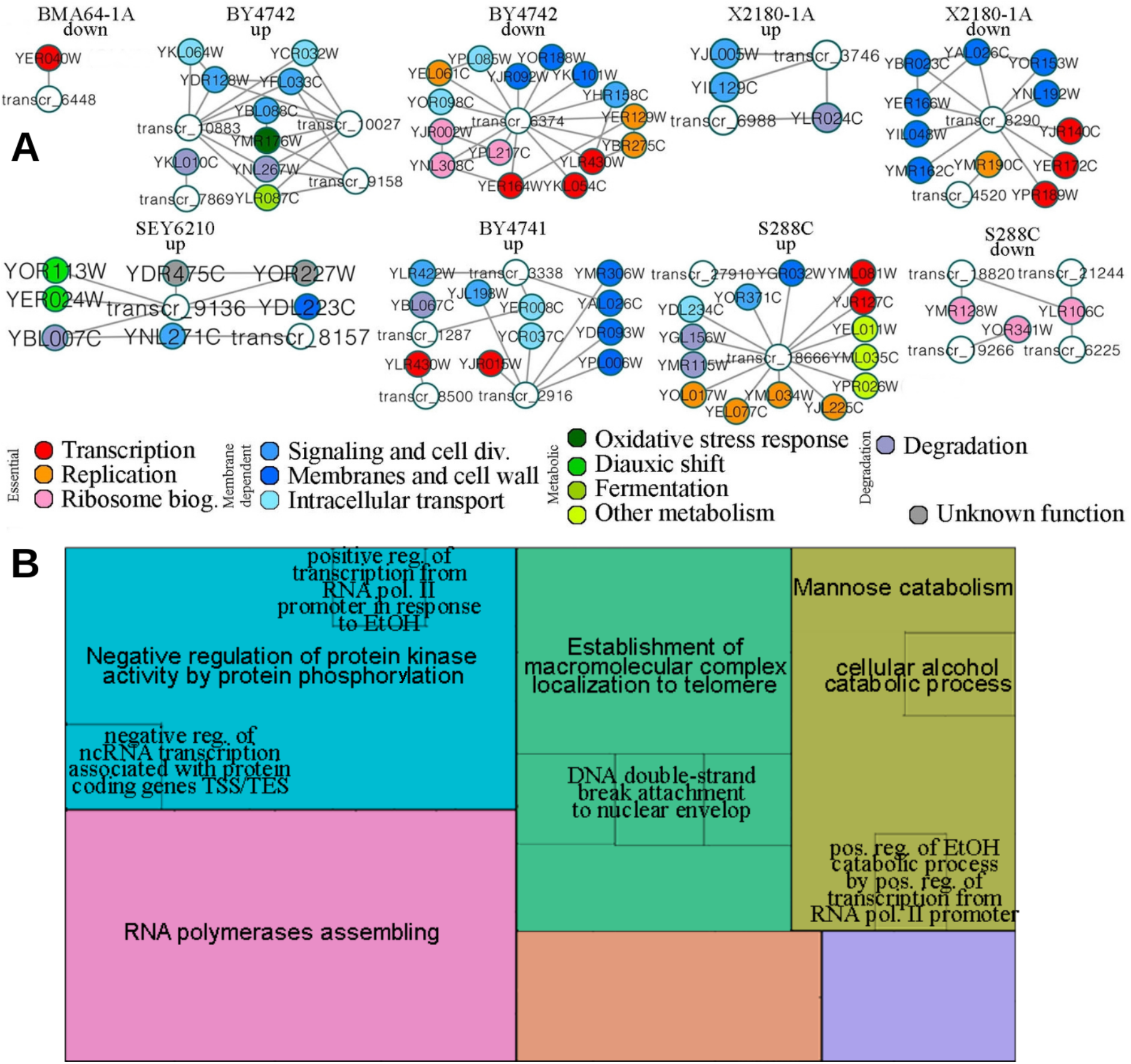
Functional annotation of lncRNAs **A:** selected subsystems using the lncRNA-propagation analysis. The node colors are related to biological functions depicted under the graphs; **B:** enriched terms of all target-proteins without considering neither strains nor differential expression and without excluding redundant proteins.

## Discussion

### EtOH stress-responsive lncRNAs are diverse and may be directly involved in the EtOH tolerance

Few lncRNAs have been found in yeasts, counting up to 18 lncRNAs for *S. cerevisiae* (Till *et al*. 2018). Here, we present the most extensive catalog of lncRNAs of 6 different *S. cerevisiae* strains; the ones were never identified before since there was no similarity to the previously lncRNAs found in yeast (Parker *et al*. 2017, 2018).

LncRNAs can act as protein-baits providing a negative impact on the targetprotein by blocking its function. However, lncRNA-scaffolders help the protein complex organization positively or negatively (Wang and Chang 2011). The LNCPI here predicted fits the number of edges (ncRNA-protein interactions) previously reported (Panni *et al*. 2017), as well as the number of target proteins (~8.1 proteins per transcript); in fact, one small RNA (100 nts) can trap around 5-20 proteins (Chujo *et al*. 2016).

We used the guilt-by-association and the information flow throughout LNCPI (the lncRNA-propagation) to assess the lncRNA functions. Overall, the EtOH tolerance responsive lncRNAs in both higher tolerant (HT) and lower tolerant (LT) phenotypes work on different pathways; we highlight the cell wall, cell cycle and growth, cell longevity, cell surveillance, ribosome biogenesis, intracellular transport, trehalose metabolism, transcription, and nutrients shifts pathways. Below, we discuss some exciting interactions that may be related to EtOH tolerance.

The down-regulated lncRNA transcr_63478 of BY4742 binds to the Cin8p (YEL061C), responsible for the mitotic spindle assembly and chromosome segregation (Roof *et al*. 1992). The lack of CIN8 leads to cell cycle progression delay (Straight *et al*. 1998; Mittal *et al*. 2020). Remarkably, CIN8 is down-regulated in all HT strains and not differentially expressed in most LTs (data not shown). Additionally, this lncRNA also binds to Def1p, an RNAPII degradation factor (Woudstra *et al*. 2002).

The EtOH affects yeast’s cell wall components (Aguilar-Uscanga and Francois 2003). The down-regulated lncRNA transcr_8290 of X2180-1A bind to the ATPases Dnf1p (YER166W), Neo1p (YIL048W), Dsr2p (YAL026C), Dnf3 (YMR162C), and chitin synthases Chs3p (YBR023C), and Chs1p (YNL192W). The 4 P-type ATPases mentioned transport phospholipids through a bilayer membrane (Paulusma and Oude Elferink 2005), and then, chitin synthases produce chitins for the cell wall (Ziman *et al*. 1996). Moreover, these proteins lie in the membrane, working as multi-drug transport in the responsive drug system (Golin *et al*. 2007). Altogether, we suggest that transcr_8290 of X2180-1A may act as an assembler of protein complexes related to membrane and cell wall.

The down-regulated lncRNAs transcr_18820, transcr_21244, and transcr_6225 of S288C bind to the Ecm16p (YMR128W), and Rea1p/Mdn1p (YLR106C), which are small nucleolar ribonucleoprotein (snoRNP) and ribosome biogenesis protein, respectively (Shiratori *et al*. 1999; Colley *et al*. 2000; Nissan 2002). Additionally, transcr_19266 interacts with Rpa190p (YOR341W), which is part of RNA Polymerase I (RNAPI) (Kuhn *et al*. 2007).

The up-regulated lncRNAs transcr_3338 and transcr_2916 of BY4741 bind to the flippases Drs2p (YAL026C) and Dnf2p (YDR093W), respectively. These flipases are P-type ATPases that concentrate phosphatidylserine and phosphatidylethanolamine on the cytosolic leaflet, contributing to endocytosis, intracellular transport, and cell polarity (Chen *et al*. 1999; Hua *et al*. 2002; Pomorski *et al*. 2003; Iwamoto *et al*. 2004). Transcr_2916 also binds to the two low-affinity phosphate transporters Pho90p (YJL198W) and Pho87p (YCR037C); the overexpression of these genes leads to abnormal cell cycle progression and a reduction of vegetative growth rate (Stevenson *et al*. 2001; Sopko *et al*. 2006; Yoshikawa *et al*. 2011). BY4741 is the only strain with up-regulation of these PHO genes (data not shown).

The up-regulated lncRNAs of SEY6210 seem to act mainly on “membrane dependent processes” in the cell modeling processes (such as the ones during the cell cycle). In this case, the interaction between transcr_8157 and transcr_9136 with the proteins Bni1p (YNL271C), Sla1p (YBL007C), and Hbt1p (YDL223C) indicate that these lncRNAs may work on the cell cortex modeling system, albeit the mentioned genes have different roles. Sla1p is a cytoskeletal binding protein associated with endocytosis or binds to proteins to regulate actin dynamics (Pruyne and Bretscher 2000; Howard *et al*. 2002). Bni1p and Hbt1p are responsible for cell polarization (Lee *et al*. 1999; Dittmar 2002; Pruyne *et al*. 2002, 2004; Tcheperegine *et al*. 2005; Guarente 2010).

The up-regulated lncRNAs transcr_18666 of S288C seems to work on trehalose metabolism actively. There is a positive correlation between cell viability and trehalose concentration in cells under EtOH stress, providing a protective effect to the EtOH (Mansure *et al*. 1994). The Ath1p (YPR026W), a trehalose degradation protein (Nwaka *et al*. 1996; Jules *et al*. 2004), interacts with transcr_18666. Interestingly, S288C has the lowest EtOH tolerance. Altogether, our initial hypothesis is that the transcr_18666 may negatively impact ATH1 under the severe EtOH stress.

The X2180-1A up-regulated lncRNA transcr_3746 may be acting on cellular surveillance mediated by a “membrane dependent process”. We found that the Cyr1p (YJL005W) (works on signal transduction, which is required for cAMP production (Kataoka *et al*. 1985)) also binds to transcr_3746. The cAMP is involved in cell cycle progression, sporulation, cell growth, stress response, and longevity (Casperson *et al*. 1985). Hence, we suggest that the transcr_3746 may also be related to cell longevity, growth, and proliferation.

The up-regulated lncRNAs transcr_10883, transcr_10027, and transcr_9158 of BY4742 may be directly involved in the EtOH tolerance controlling nutrient supply. These lncRNAs bind to Pik1p (YNL267W), a kinase that can rapidly restore the nutrients supply in cells under nutrient deprivation (Demmel *et al*. 2008). Interestingly, cells under EtOH stress ongoing by a variation on nutrients depletion (Tesnière *et al*. 2013).

The down-regulated lncRNA transcr_6448 of BMA64-1A had one target-protein coded by the gene YER040W (GLN3). Gln3p is a transcriptional activator of genes subject to nitrogen catabolite repression (Feller *et al*. 2013). The deletion of GLN3 boosts the yeast to the branched-chain alcohols tolerance (Kuroda *et al*. 2019). The finding mentioned fits our hypothesis since BMA64-1A is the only strain with the downregulation of GLN3 (data not shown) and presents the highest EtOH tolerance here analyzed. In this case, we suggest that the transcr_6448 may act as a sort of Gln3p repressor under the severe EtOH stress by an unknown negative-feedback loop. Altogether, we hypothesize that the highest EtOH tolerance observed for BMA64-1A may be a by-product of the negative GLN3 regulation by the transcr_6448 under stress.

Plenty of lncRNA’s target-proteins are related to transcriptional mechanisms (“RNA polymerase assembling” and “negative regulation of protein kinase activity by protein phosphorylation”). The proteins related to the “RNA polymerase assembling” mechanism are related to the “transcription by RNA polymerase II”. LncRNAs can contribute to a balance of transcription/degradation rate (Timmers and Tora 2018). Therefore, we hypothesize that lncRNA-RNAPII interactions might be acting as signaling molecules to counterbalance the transcription/degradation rate of mRNAs.

Altogether, network analysis provided hints about the role of many EtOH stress-responsive lncRNA. These ncRNAs seem to act as baits, backbones, or adapter molecules to maintain protein complexes essentials to improve yeast capacity to endure the severe EtOH stress. Finally, our findings indicate that yeasts’ lncRNAs under a severe EtOH stress seem to interconnect a diversity of modules to surpass hurdles imposed by this stressor.

